# Musculoskeletal modelling of sprawling and parasagittal forelimbs provides insight into synapsid postural transition

**DOI:** 10.1101/2021.06.15.448564

**Authors:** Robert J. Brocklehurst, Philip Fahn-Lai, Sophie Regnault, Stephanie E. Pierce

## Abstract

The sprawling–parasagittal postural shift was a major transition during synapsid evolution and is considered key to mammalian ecological diversity. Despite a good fossil record, debate remains over when the shift to parasagittal posture occurred, primarily due to limited comparative biomechanical data on extant species. Here, we built forelimb musculoskeletal models of three extant taxa that bracket the sprawling–parasagittal transition: a tegu lizard, an echidna, and an opossum. We measured shoulder joint range of motion (ROM) about all three degrees of rotational freedom and characterized shoulder muscle moment arms (MMAs) across the entire pose space. Our results show that both the opossum and the tegu had high shoulder joint ROM, and both were substantially higher than the echidna. However, the opossum occupied a distinct region of pose space characterized by high humeral retraction angles. There are clear interspecific differences in MMAs related to posture, with the sprawling tegu and echidna emphasizing humeral depression, and the parasagittal opossum emphasizing humeral elevation. There are also notable differences between our sprawling taxa, with the echidna possessing much greater moment arms for humeral pronation than the tegu. We demonstrate clear functional variation between locomotor grades and use these data to hypothesize major shifts in forelimb function and posture along the mammalian stem.

## Introduction

Extant therian mammals (marsupials and placentals) can be distinguished from other land-living quadrupeds by their derived parasagittal gait, with the limbs adducted close to the body midline and the limb joints aligned in a single plane [1,2]. This contrasts with the sprawling postures plesiomorphic for tetrapods – still typified by extant non-avian reptiles and amphibians – which are characterized by abducted limbs and multiaxial joints [1,3,4]. A postural shift from sprawling to parasagittal is heralded as a major transition during mammalian evolution and has been hypothesized to confer several adaptive advantages [1,5]. As well as possessing distinctive limb postures, extant therians occupy a wide array of niches and the forelimbs have been exapted and transformed to serve diverse ecological functions [6]. Both the parasagittal posture and increased functional disparity of therian forelimbs have been attributed to their distinct shoulder morphology: the shoulder girdle is reduced and highly mobile, the ventrally facing glenoid means the limbs project ventrally rather than laterally, and the “ball-and-socket” glenohumeral joint is hypothesized to permit a wider range of motion [7].

The parasagittal posture and functionally diverse forelimbs of therians contrast with the sprawled posture and functionally constrained forelimbs of their earliest ancestors among the non-mammalian synapsids [8,9]. Synapsids possess a good fossil record that documents the origins of key mammalian traits [5], yet there remains considerable disagreement over when major functional changes in forelimb use and posture occurred during the evolution of mammals. This is primarily due to differing functional interpretations of preserved skeletal anatomy and reconstructed soft-tissue [5,8,10,11], a problem exacerbated by limited biomechanical data from extant taxa, particularly on how bones and soft tissues interact to produce forelimb movement. Due to the disparity in their forelimbs, parasagittal and sprawling taxa are often examined independently of one another, with few studies explicitly comparing sprawling vs. parasagittal locomotion within the same biomechanical framework [12,13]. Therefore, we currently lack comparative functional morphology data from extant taxa that may allow us to better interpret functional evolution along the mammalian stem lineage.

To address this data gap, we built forelimb musculoskeletal models of three extant taxa that functionally span the sprawling–parasagittal transition: the Argentine black and white tegu (*Salvator merianae*), a representative squamate reptile with a classical sprawling gait, combining humeral long-axis rotation and limb retraction (flexion in a horizontal plane) [4,12]; the short-beaked echidna (*Tachyglossus aculeatus*), a monotreme mammal with an “upright-sprawling” gait, in which limb movement is primarily driven by humeral long-axis rotation (and rolling of the trunk) [14–16]; and the Virginia opossum (*Didelphis virginiana*), a therian mammal with a parasagittal gait, that mainly uses limb elevation (flexion in a dorsoventral plane) [17,18]. At the glenohumeral joint, we measured three-dimensional osteological range of motion (ROM) and muscle moment arms (MMAs), two popular metrics among functional morphologists for comparing locomotor behaviour across fossil taxa and their extant analogues [19]. ROM data are commonly used to define the limits of possible poses that an animal might use in life, and have been used to reconstruct locomotor behaviour in fossil species [20,21]. MMAs represent the leverage of muscles’ around a joint and have been used to infer adaptations of the musculoskeletal system towards specific types of locomotor behaviours in extant and fossil taxa [16,19,22]. Although ROM-MMA analyses are typically performed using a ‘single axis’ approach, MMAs covary with all rotational degrees of freedom [23]. As the sprawling–parasagittal postural transition involved major reorganization of the musculoskeletal system, understanding how joint and muscle function changed across the transition requires exploring ROM and MMAs within a 3D pose space.

Here we explored broad-scale relationships between shoulder joint morphology, mobility, and muscle function across a large range of feasible limb poses and postures by modelling multiple degrees of freedom simultaneously [24]. Based on previous studies of *in vivo* locomotion kinematics, osteological ROM, and interspecific differences in joint and limb morphology [4,12,14–18,25], we predicted that shoulder joint ROM would be lowest in the echidna, intermediate in the tegu and highest in the opossum. We also predicted that MMAs about the glenohumeral joint would be highest in those specific degrees of freedom most important for locomotion: long-axis rotation and retraction in the tegu; long-axis rotation in the echidna; and elevation in the opossum [12,14,16–18]. Our results show that there is a complex relationship between posture, morphology, and joint and muscle function, but also reveal key similarities and differences between both sprawling and parasagittal taxa and between different types of sprawlers. We use these data, in combination with anatomical transformations in the fossil record, to hypothesize when major changes in forelimb function and posture occurred during synapsid evolution.

## Materials and Methods

### Shoulder Musculoskeletal Geometry

All specimens used here were digitized as part of previous studies [13,16,26]. Briefly, specimens were contrast stained with Lugol’s iodine and micro-CT scanned (for specimen details and scan parameters, see table S1) to capture both hard- and soft-tissue anatomy. Tomographic images were segmented using Mimics version 19 (Materialise NV, Leuven, Belgium) to extract musculoskeletal geometry. Muscle insertion and origin sites were painted onto the forelimb bones in Mudbox (versions 2019, 2020, Autodesk Inc., San Rafael, CA, USA) and extracted as individual meshes. To facilitate creation and alignment of anatomical and joint coordinate systems [27], geometric primitives – planes, spheres, cylinders – and convex hulls were fit to the joint articular surfaces of the bones (i.e., glenoid, proximal and distal humerus) in Meshmixer (Autodesk Inc., San Rafael, CA, USA). 3D models of the bones, muscles, muscle insertion sites and joint primitives were imported into Maya (versions 2019, 2020, Autodesk Inc., San Rafael, CA, USA) as .obj files for further analysis and construction of the biomechanical models. In Maya, centroids of the muscle attachment meshes were calculated using the “vertAvg” tool from the XROMM toolbox (available at https://bitbucket.org/xromm/xromm_mayatools/wiki/Home). For muscles with more extensive soft-tissue attachments that could not be easily mapped onto bones, e.g. the mm. latissimus dorsi and pectoralis, origin and insertion points were placed manually based on 3D muscle geometry [13,16].

### Model Assembly

To facilitate comparisons across animals with very different habitual postures, all models were initially positioned in a global ‘reference pose’, at which all joint angles equaled zero (figure 1). This reference pose generally follows previous studies [28] and is constructed as follows. The pectoral girdle was anatomically aligned with the global axes and the joint coordinate system for the glenoid was oriented in the same way (see figure S1). The humerus was then aligned with the glenoid such that the humerus’ long axis pointed laterally, parallel to the global X-axis, and the distal humerus parallel to the ground (figure 1A). In this configuration, glenohumeral joint rotation about the dorsoventrally oriented Z-axis represents humeral retraction and protraction; rotation around the craniocaudally oriented Y-axis represents humeral depression and elevation (adduction and abduction); and rotation about the mediolaterally oriented X-axis represents humeral long-axis rotation (pronation and supination) (figure 1A). Protraction-retraction and elevation-depression are akin to longitude and latitude on a sphere, and long-axis rotation denotes the heading [29]. The elbow joint was oriented so that the global zero was an extended pose, with the long axis of the antebrachium parallel to the horizontal plane (figure 1, figure S1).

**Figure 1.**
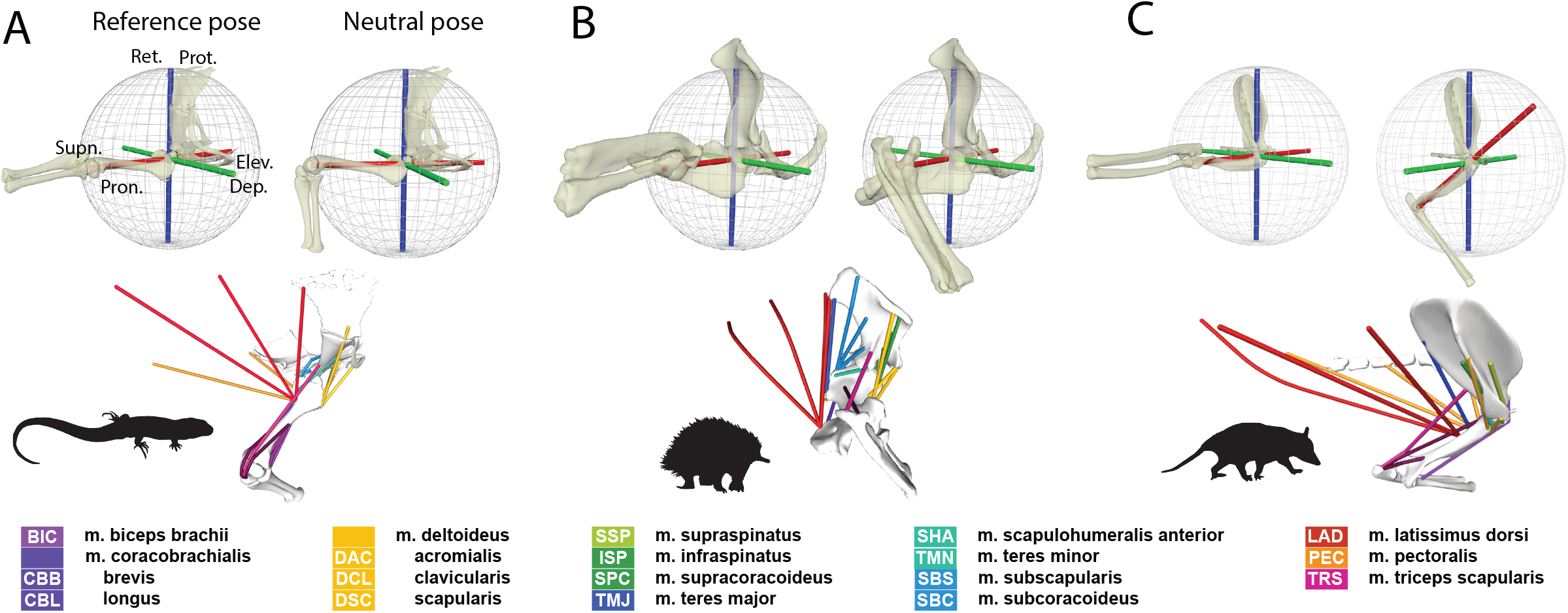
Musculoskeletal models of the forelimb in the (A) tegu, (B) echidna and (C) opossum. Top row: global reference pose and anatomically informed neutral poses. The blue Z-axes are (-) retraction and (+) protraction, the green Y-axes are (-) depression and (+) elevation and red X-axes are long-axis rotation or (-) pronation and (+) supination. Bottom row: OpenSim models showing locations of the studied muscles with homologous muscles color coded as detailed in the legend. See SI for further information on model construction.

Separate from the global reference pose, we also set up an anatomically informed ‘neutral pose’, based on joint anatomy, which was distinct for each animal (figure 1A). Based on comparison with experimental data [12,14,18], each neutral pose was considered representative of a pose the animals might use in life and was used as the ‘starting pose’ during the ROM analysis. However, all angles were measured relative to the global reference pose. To generate the neutral pose, local anatomical coordinate systems were created and aligned based on the articular surface morphology of the shoulder joint (glenoid and humeral head) (figure S2). The proximal and distal anatomical glenohumeral axes were then superimposed, and joint spacing adjusted by translating the humerus along the anatomical X-axis, oriented normal to the glenoid. For all three taxa, minimum joint spacing – assessed by extruding the humeral head until it contacted the glenoid – was less than 1% humeral length. For the elbow, we used the same global axis system, but the joint was flexed at 90° for the anatomically informed starting pose.

### Measuring and Analyzing Range of Motion (ROM)

Osteological range of motion (ROM) was measured in Maya, by modifying published automated methods [24]. Starting from the anatomically aligned neutral pose, the glenohumeral joint was moved through a series of rotation combinations – protraction-retraction angle ±90°, elevation-depression angle ±90°, long-axis rotation angle ±90°, in that rotation order – with increments of 10° (totaling 6,859 poses). We chose to only sample within ±90° of the starting pose rather than sampling all possible poses because many regions of pose space, while osteologically viable, are not biologically realistic [16,24]. Surveying published *in vivo* data, animals rarely move their limbs through joint excursions >90° during quadrupedal locomotion [4,12,18,28], and as the anatomical neutral pose should generally fall within *in vivo* range of motion, we would expect these ROM envelopes to capture most poses used in life.

For each combination of joint angles, poses were deemed viable if there was both no bone-to-bone interpenetration, and the joints were still articulated i.e. if joint surfaces were deemed to overlap (figure S3). For each species, the complete list of viable poses was exported from Maya as a .csv file. These files were then imported into R for further analysis and visualization. We performed a cosine-correction of our 3D pose space [29], as this allows more accurate inter-model comparisons. We compared overall joint mobility by calculating the volume of alpha shapes or “concave hulls” fitted to the viable poses for each taxon in cosine-corrected pose space using an alpha value of 10 [24,29].

### Measuring and Analyzing Muscle Moment Arms (MMA)

Information from the Maya scene – bone meshes, joint axis orientation and position, and shoulder muscle insertion and origin position – was converted into an OpenSim (version 4.1) model using a set of custom-written Python and R scripts. In OpenSim [30], additional refinements to the models were made, such as the addition of wrapping surfaces and via points to constrain the muscle lines of action to anatomically realistic paths [22]. For each species, all viable glenohumeral joint poses were exported from Maya and converted to motion (.mot) files. These motion files were imported into OpenSim, and MMAs calculated about each rotational degree of freedom for each pose. This novel approach characterizes MMAs across all feasible glenohumeral joint angles, providing a more comprehensive view of muscle function in relation to limb pose and posture. For all glenohumeral joint poses tested, the elbow remained flexed at 90°.

The MMA data were imported into R for analysis and visualization. MMAs were normalized to specimen size by scaling to humeral volume^1/3^. At each viable pose for each degree of freedom, we calculated the sum of the normalized moment arms in each direction (positive and negative), to compare across species. When calculating the sums, mean MMA values were calculated for muscles with multiple modelled origins and insertions to avoid overrepresentation skewing the summed MMA results. We took means for mm. biceps, mm. coracobrachialis, m. subscapularis (echidna only), m. infraspinatus (echidna only) and m. latissimus dorsi [16]. We also took mean MMAs for the middle plus caudal parts of the m. pectoralis, and for the m. deltoideus scapularis plus m. deltoideus acromialis based on similar electrical activation patterns [12,16]. We plotted the distributions of summed normalized muscle moment arms across pose space and compared them between our three taxa. Due to the large number of datapoints, we also summarized the distributions of both summed MMA and individual muscle MMA across pose space using boxplots and density curves.

## Results

### ROM Analyses

The opossum had the greatest overall glenohumeral joint ROM, followed closely by the tegu (alpha hull volumes 1,914,210 degrees^3^ and 1,300,807 degrees^3^ respectively), and both were orders of magnitude greater than the echidna (alpha hull volume 89,339 degrees^3^). Looking at the shape of the 3D ROM envelopes, the echidna’s ROM envelope is narrowest along the protraction-retraction axis, and widest along the elevation-depression axis (figure 2), consistent with prior work [16]. ROM in the tegu and the opossum is generally more evenly distributed in all three dimensions, although their ROM envelope positions within pose space are different due to differences in the neutral pose and joint anatomy (figure 2). The sampled poses in the opossum occur at higher retraction and depression angles than those of the tegu (figure 2), reflecting the differences in habitual limb posture (parasagittal vs. sprawled) and glenoid orientation (ventral vs. lateral).

**Figure 2.**
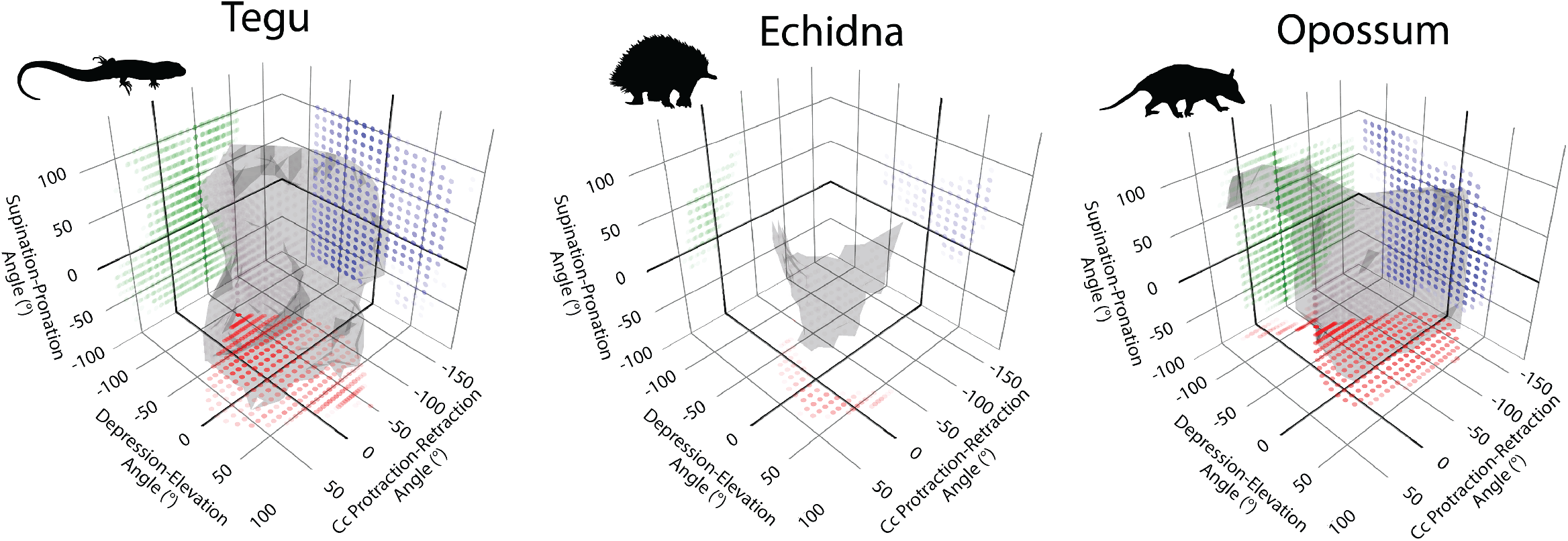
Range of motion (ROM) envelopes for the glenohumeral joint in the tegu, echidna and opossum. ROM envelopes are plotted in cosine-corrected Euler space. Projected points are color coded according to the axis of rotation normal to the plane (red on the ZY plane, green on the ZX plane, blue on the XY plane). For interactive version of this plot see supplementary shiny app.

### MMA Analyses

#### Summed MMA: Global patterns

Looking at the distributions of summed normalized MMAs across pose space for each species (figure 3A) and their numerical summaries (figure 3B), the echidna has the highest median value for summed humeral retraction MMAs (figure 3B), but the opossum has the highest peak retraction values (figure 3A,B). These high retraction MMA values in the opossum occur when the humerus is at low retraction angles (> -50°) (figure 3A). Median values for humeral protraction are greater in the tegu and echidna than in the opossum (figure 3B), but the opossum and tegu have higher peak values. These peak values of protraction occur in the tegu either when the humerus is elevated and pronated or depressed and supinated (figure 3A), and at extreme retraction angles in the opossum (figure 3A).

**Figure 3.**
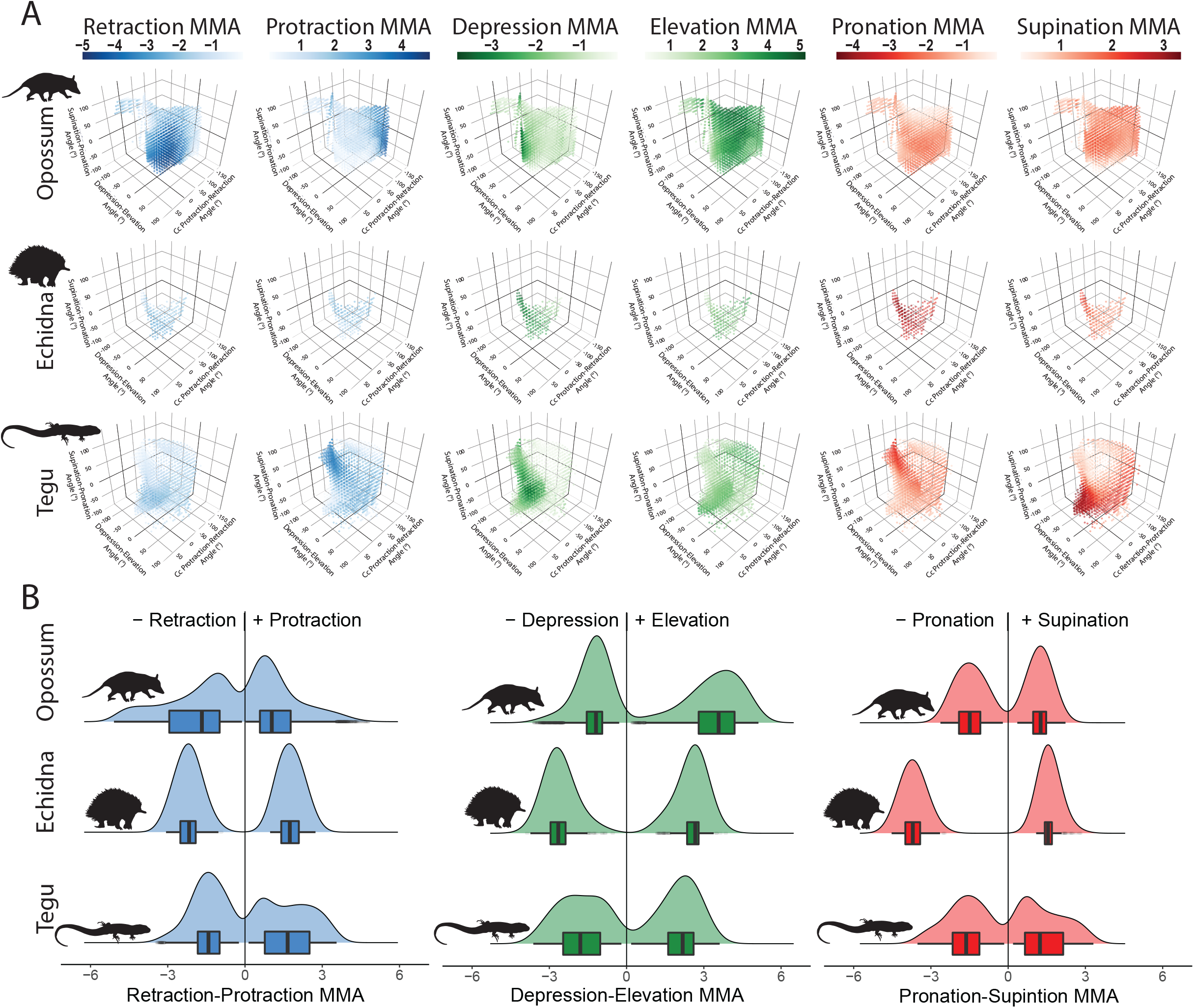
Muscle moment arms (MMAs) across pose space in the tegu, echidna and opossum. (A) Summed normalized MMAs at each glenohumeral joint angle. (B) Summary distribution and boxplots showing medians for summed normalized muscle moment arms. For interactive version of panel (A) see supplementary shiny app.

For humeral depression, the opossum has the lowest median MMA values, and the echidna has the highest; however, peak values for depression are similar in the echidna and the tegu (figure 3B). In both the tegu and echidna, humeral depression MMAs are greatest at more depressed joint angles (figure 3). Peak and median MMA values for elevation are lowest in the tegu, intermediate in the echidna, and greatest in the opossum (figure 3B). The opossum has generally high values for summed elevation MMAs, but they are greatest at high retraction angles and with the humerus depressed below the horizontal (figure 3A).

Both median and peak MMA values for humeral pronation are substantially higher in the echidna than in either the tegu or the opossum (figure 3B), and they are distributed evenly across the echidna’s pose space (figure 3A). Median MMA values for supination are greatest in the echidna but peak values are highest in the tegu (figure 3B). These peak values occur when the tegu humerus is pronated (figure 3A).

#### Individual MMA: Muscle function

The observed similarities and differences in overall summed MMAs are the cumulative result of the actions of individual muscles (figure 4, supplementary shiny app). The mm. biceps brachii (BIC) has similar actions depressing the humerus in all three taxa and pronating it in the tegu and opossum. The m. coracobrachialis (CB) depresses the humerus in the echidna and tegu but depresses and elevates in the opossum. This muscle also has larger pronation MMAs in the tegu and echidna. The acromial and scapular portions of the m. deltoideus (DAC + DSC) have similar functions in all three taxa, elevating, supinating and protracting the humerus, whereas the clavicular portion (DCL) behaves differently; in the tegu and echidna it protracts and supinates the humerus, but in the opossum it pronates it.

**Figure 4.**
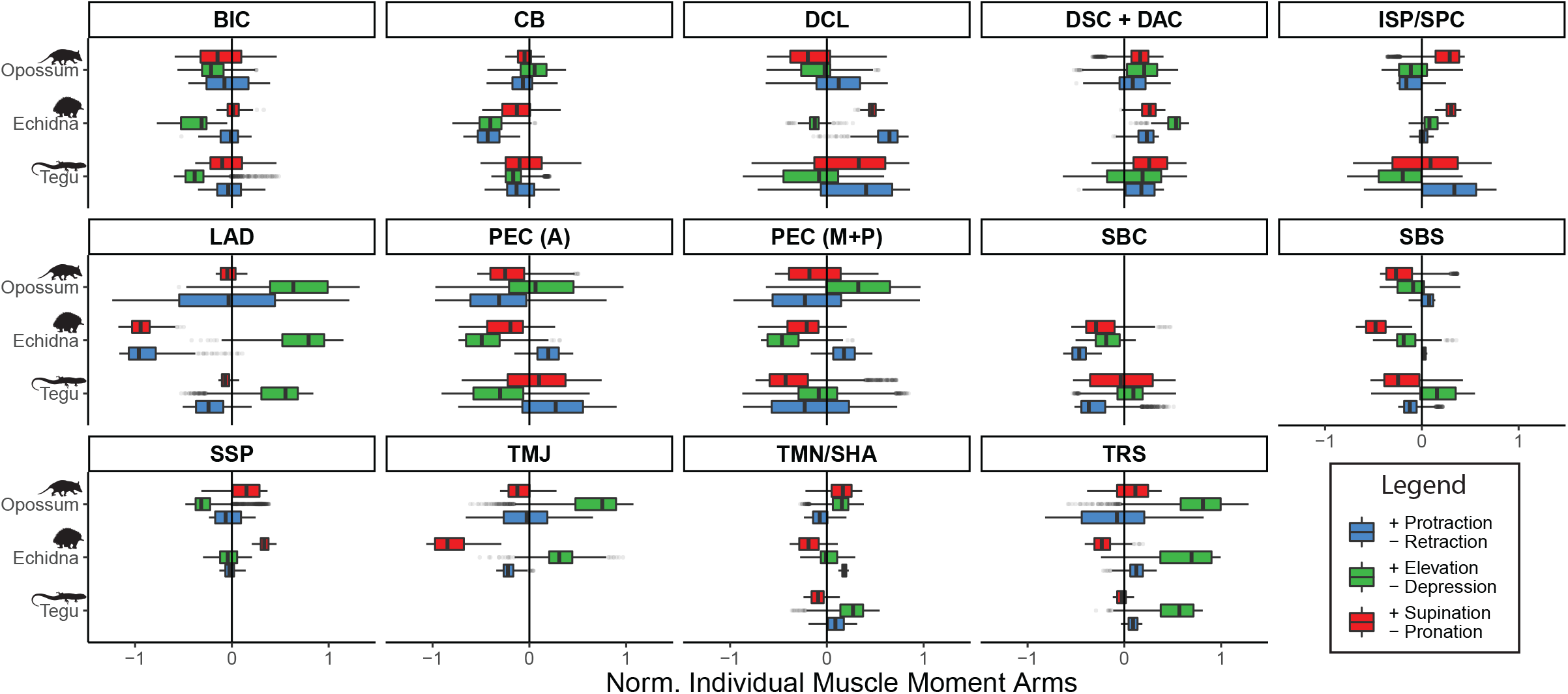
Boxplots showing median values and distributions of normalized muscle moment arms (MMAs) for the different muscles modelled in the tegu, echidna and opossum. Homologous muscles are indicated by a forward slash. Aggregated muscles are indicated by a plus. See Figure 1 for muscle name abbreviations. For interactive plots of MMAs across pose space see supplementary shiny app.

The m. infraspinatus (ISP) of the opossum has MMAs for supination, depression, and retraction, but in the echidna this muscle elevates the humerus. The homologous muscle in the tegu, the m. supracoracoideus (SPC), also depresses the humerus, but protracts rather than retracts (figure 4). The m. supraspinatus (SSP) is only present in the opossum and echidna and supinates and retracts in both, but also depresses in the opossum. The m. subcoracoideus (SBC) is only present in the tegu and echidna and acts to retract the humerus in both taxa, while also elevating in the tegu, but depressing and pronating in the echidna (figure 4).

The m. teres minor (TMN) of the opossum and echidna is homologous with the tegu m. scapulohumeralis anterior (SHA) [13]. In the tegu and opossum, this muscle acts to elevate the humerus but it elevates and depresses in the echidna. In the tegu and echidna, this muscle also pronates and protracts the humerus but has the opposite actions (retraction and supination) in the opossum. The m. teres major (TMJ) is present only in the opossum and echidna. It both retracts and protracts in the opossum, but only retracts in the echidna (figure 4). In both taxa this muscle elevates and pronates the humerus, but the echidna has larger pronation MMAs and the opossum has larger elevation MMAs (Figure 4).

In all three taxa studied, the m. latissimus dorsi (LAT) elevates and pronates the humerus, but whereas in the tegu and echidna this muscle strictly retracts, in the opossum it both protracts and retracts.

M. latissimus retraction and pronation MMAs are particularly high in the echidna, and elevation MMAs are highest in the echidna and opossum. The function of the m. pectoralis (PEC) differs between taxa. In the opossum, both anterior (PEC A) and more posterior parts (PEC M+P) elevate, pronate, and retract the humerus, while in the echidna, both portions serve to protract, depress and pronate the humerus. The tegu possesses different MMAs in different parts of the muscle; anteriorly it supinates, depresses, and protracts the humerus, but posteriorly it pronates and retracts, with lower depression and greater elevation MMAs (figure 4). The m. triceps scapularis (TRS) has high MMAs for humeral elevation in all three taxa. In the tegu and echidna this muscle has protraction and pronation MMAs but supinates the humerus more in the opossum and has higher MMAs for both retraction and protraction (figure 4).

## Discussion

### Therian ball-and-socket shoulder joint is not uniquely mobile

The mobile ball-and-socket shoulder (glenohumeral) joint of therian mammals has been suggested to underpin their extant ecological diversity, possessing a wide ROM envelope and allowing the forelimb to perform a greater range of movements and associated functions [6,7]. Our results confirm that the shoulder joint in the opossum is indeed highly mobile as predicted, especially compared to its immediate sister taxon in this study, the sprawling monotreme echidna (figure 2)(also see [14,16]). However, the tegu also had high shoulder joint ROM, and both the opossum and tegu had considerably higher ROM (orders of magnitude) than the echidna (figure 2). In particular, the tegu and opossum had much higher ranges of motion for humeral protraction-retraction and long-axis rotation than the echidna. This result indicates that the ‘hemi-sellar’ glenohumeral joint morphology of the tegu and echidna function differently, and that such joints are not inherently restrictive to motion [25]. Therefore, when compared to more ‘classical’ sprawling outgroups (e.g., lepidosaurs and crocodilians) with greater ROM at the shoulder joint than monotremes [19,20,28], the high ROM permitted by the ball-and-socket shoulder joint in therians seems less extreme, and a less compelling explanation for therian forelimb diversity. However, the therian shoulder joint does operate in a fundamentally different region of pose space (figure 2) – at generally higher retraction and depression angles [17]. This correlates with differences in glenoid orientation [7,11], and leads to major differences in limb loading regimes [2,3].

### Shoulder muscle elevation leverage key to parasagittal posture

Based on experimental studies of joint kinematics and muscle activation patterns [17,18], we predicted the opossum would have the greatest summed MMAs for humeral elevation, as under our coordinate system humeral elevation at high retraction angles corresponds to flexion in the parasagittal plane [18]. The opossum did indeed have the highest MMAs for humeral elevation out of the three taxa studied here (figure 3). Increased MMAs for forelimb elevation – particularly at high glenohumeral retraction angles – may therefore be an important part of the reorganization of the musculoskeletal system over the course of mammalian evolution.

The increased MMA for elevation in the opossum is due to several factors. The m. teres major is a muscle that imparts a large elevation MMA (figure 4), and although it has been lost in some leipdosaurs, it has been reconstructed as plesiomorphic for Amniota and is present in mammals, crocodilians and turtles [31]. However, comparing our two mammals, the m. teres major still has larger elevation MMAs in the parasagittal opossum than in the sprawling echidna (figure 4). There are also several muscles that generally act as elevators in all three taxa but have greatest MMAs in the opossum (e.g., m. triceps scapularis), or which have different functions between taxa – acting as elevators in the opossum but depressors in the tegu and echidna (e.g., m. coracobrachialis, m. pectoralis) (figure 4).

Of the muscles whose elevation-depression actions differ between taxa, the m. pectoralis is of key interest as the muscle appears to change its function across pose space (see supplementary app). In both the tegu and opossum, which share large regions of pose space (figure 2), the m. pectoralis showed a pattern whereby its MMA trends from depression to elevation with increasing humeral retraction angles (figure S4; also see supplementary shiny app.). While the m. pectoralis is an important humeral retractor and antigravity ‘adductor’ muscle in sprawling tetrapods [16,32], it helps drive locomotion via humeral elevation during the power stroke of stance in parasagittal therians [18]. Therefore, our results imply that during the sprawling–parasagittal transition, changes in muscle function may have come about purely through increasing humeral retraction angles via reorientation of the glenohumeral joint.

### Echidna and tegu sprawl in different ways: spin vs. swing

The tegu and echidna are both classed as ‘sprawlers’ based on their abducted limbs and low humeral retraction angles [12,14,17]. They shared some similarities in MMAs consistent with this e.g., high MMAs for humeral depression (figure 3). In sprawling animals, humeral depression corresponds to the adduction action of support musculature resisting abduction moments generated by laterally placed ground reaction forces [3,12]. However, there were also several key differences between the tegu and the echidna. The echidna had the greatest summed MMAs for humeral pronation (figure 3) as expected based on its reliance on humeral long-axis rotation during propulsion [14,16]. Many of the echidna’s individual muscles possessed higher MMAs for pronation than their homologues in the tegu (figure 4), but particularly the m. latissimus dorsi, which has a derived attachment site on the distal humeral entepicondyle [26]. The echidna also possesses muscles absent in the tegu; of these, the m. teres major has a substantial pronation MMA (figure 4).

Contrary to our predictions, the tegu did not have greater MMAs than the echidna for humeral retraction (figures 3, 4), despite the prominent role of retraction in the classical sprawled gait [4,12]. In fact, the tegu had lower MMAs than the echidna in almost all degrees of freedom (figure 3). Although the echidna possesses extra muscles, many of the homologous muscles present in both taxa also have lower median MMAs in the tegu than the echidna (Figure 4). This is also true if we only compare poses which are viable in both taxa (figure S5). Compared to the echidna, many muscles in the tegu insert more proximally on the humerus, and the humerus itself is much more gracile [13,26]; this brings the muscle insertion sites closer to the axes of rotation, reducing MMAs but increasing the arc of the limb.

Given the difference in shoulder joint mobility between these two taxa (figure 2), and the known trade-off between MMA and a muscle’s working range (for a given fibre length [33]), we interpret these results in terms of balancing different aspects of muscle function. Studies of forelimb muscle properties in both the echidna and tegu show that there is little specialization in terms of working range or force production, and generally the muscles have low architectural disparity [13,26]. The high MMA values seen in the echidna indicate adaptation for production of higher joint moments, with the reduced muscle working range rendered moot by the restrictive nature of the glenoid. On the other hand, the tegu forelimb has generally lower MMAs and high glenohumeral ROM, which may represent adaptation for force production across a high muscle working range within the constraints of unspecialized muscles with average fibre lengths.

### Major functional shifts in synapsid forelimb evolution

Our biomechanical framework highlights several similarities and differences between the tegu, echidna, and opossum that can provide a powerful means to interpret postural shifts in the fossil record. Below, we use the results from our musculoskeletal models to predict how shoulder function may have changed across the sprawling–parasagittal transition in synapsids and put forward hypotheses on the evolution of joint and muscle function at major ancestral nodes along the mammalian stem (figure 5).

**Figure 5.**
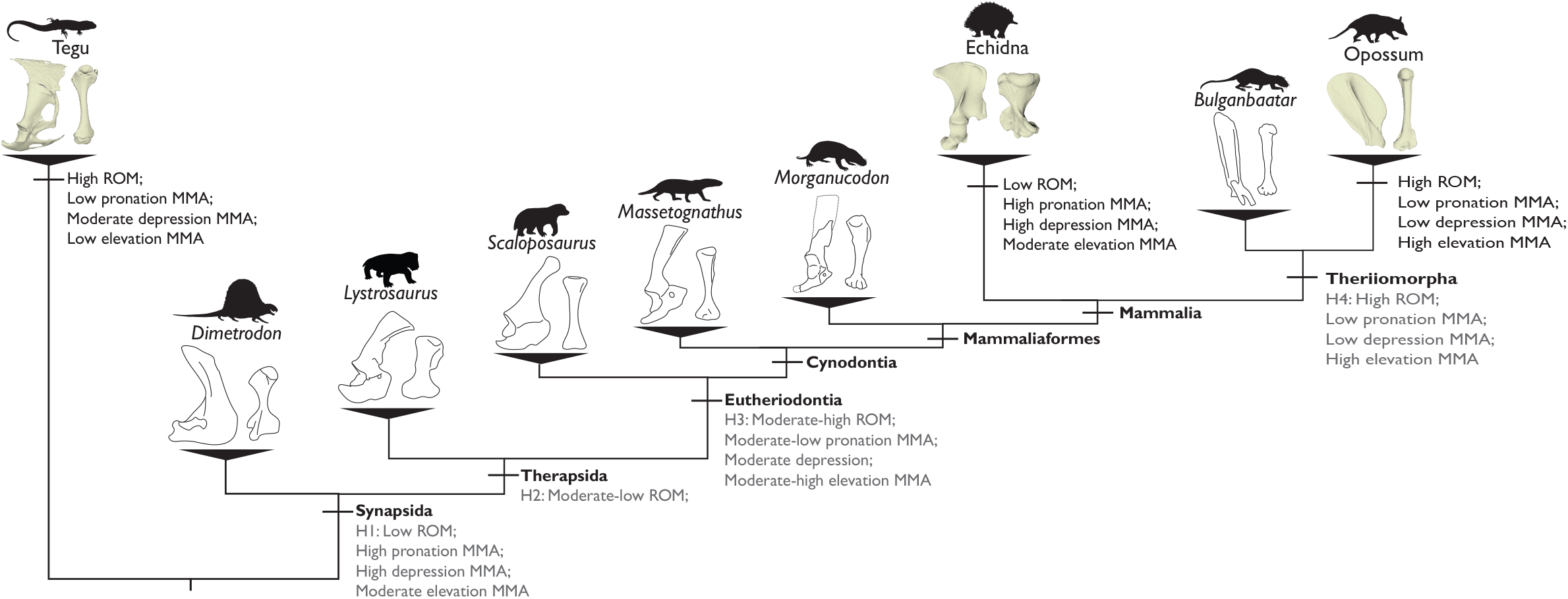
Evolution of synapsid forelimb function plotted across a simplified phylogeny, showing scapulae and humeri for representative taxa. Black annotations are results from the present study; grey annotations are hypothesized changes in forelimb ROM and MMAs based on anatomical transformations observed in fossil synapsids. Silhouettes taken from Phylopic or traced from artwork by Bogdanov (*Dimetrodon*) and Ugueto (*Scaloposaurus* and *Massetognathus*).

The most basal grade of fossil synapsids, the ‘pelycosaurs’, possessed a screw-shaped glenoid, which is proposed to greatly constrain the motion of the humerus to mainly long-axis rotation [8,11,34]. The humerus of ‘pelycosaurs’ also featured expanded epiphyses for muscle attachment [8] (figure 5). Although historically compared to lepidosaurs [8,34], ‘pelycosaurs’ share aspects of their forelimb functional morphology with the echidna; restricted glenohumeral ROM, emphasis on humeral long-axis rotation movements, and expanded epiphyses [16,34,35]. Therefore, we might expect ‘pelycosaurs’ to exhibit large MMAs, particularly for pronation, but not as extreme as those of the echidna which possess additional derived features, e.g., distal insertion of the m. latissimus (figures 3, 4). ‘Pelycosaurs’ are reconstructed with a m. teres major [10,31], and likely had moderate elevation MMAs, as this muscle had a lower elevation MMA in the sprawling echidna than in the opossum (figure 4). We also predict high depression MMAs, as the large epiphyses increase moment arms of the m. pectoralis, which functions as a depressor in sprawlers like ‘pelycosaurs’ (figure 4).

With the origin of therapsids, the glenoid joint lost its craniocaudal screw-shape, instead becoming a dorsoventrally-oriented notch [5]. However, most early-branching and large bodied therapsids, including dinocephalians and anomodonts, retained laterally expanded humeral epiphyses [9,35] (figure 5). While ROM is predicted to be higher than in ‘pelycosaurs’ because of changes to the glenoid joint, the expanded humeral epiphyses would still restrict ROM to some extent, as demonstrated in the echidna (figure 2). In more derived therapsids, and with the origin of non-mammalian cynodonts, there is a retention of the glenoid notch but a distinct shift towards more slender, gracile humeri [9–11,36] (figure 5). This combination of traits is hypothesized to increase shoulder joint mobility, particularly in protraction and retraction [10,36] as demonstrated in the tegu (figure 2), and result in reduced MMAs and a greater balance of leverage between different degrees of freedom. The glenoid also faced more caudally [10,36], indicating the forelimbs may have operated at higher shoulder joint retraction angles. Our data suggest that higher retraction angles increase MMAs for elevation and reduce MMA for depression by changing the actions of certain muscles e.g., the m. pectoralis (figures 4 and S4; also see supplementary shiny app.).

Although glenoid morphology remains notch-like in mammaliaforms [7], there is some diversity in the ‘openness’ of the facet [37], and considerable variation in humeral shape [35]. Indeed Mesozoic mammaliaforms explored several new niches not seen in non-mammalian cynodonts [37,38], additional evidence that forelimb-use and ROM is not necessarily inhibited by the possession of hemi-sellar joints. Further transformation of the shoulder joint in more crownward taxa reflects a trend towards a more parasagittal posture [7] (figure 5). Within stem therians, we see the glenoid reorient to face ventrally as opposed to laterally, the shape changes to the spheroidal socket of modern therians, and the coracoid contribution to the joint is lost [7]. These traits are considered hallmarks of the mammalian forelimb and are hypothesized to indicate the origin of forelimb parasagittal posture and gait [39,7].

## Conclusion

The origin of parasagittal posture and gait was a major event in the evolution of mammals, associated with fundamental reorganization of the musculoskeletal system, and expansion of forelimb functional disparity and ecological diversity. Here we provide a biomechanical framework to quantify how sprawling and parasagittal forelimb postures and morphologies affect glenohumeral joint mobility, muscle function and ultimately potential for forelimb use. Our data show that the parasagittal opossum had the highest shoulder joint ROM, consistent with its ball-and-socket joint morphology. However, the sprawling tegu also had high ROM, substantially higher than the echidna, indicating that glenoid morphology is not the sole determinant of shoulder mobility. Furthermore, our MMA results reveal important similarities and differences in muscle function both between and within postural grades. Our sprawling taxa both emphasized humeral depression MMAs, which is necessary to resist the elevating (abducting) moments generated by the more horizontally directed ground reaction forces. This contrasted with the opossum, in which humeral elevation MMAs are emphasized. At the high shoulder retraction angles observed in the opossum, elevation acts to produce flexion in the parasagittal plane – the characteristic movement of therian locomotion. There were also obvious differences between our sprawlers; the echidna strongly emphasizes pronation, while the tegu has a more equal distribution of MMA in multiple degrees of freedom. This dataset gives us the unique opportunity to better conceptualize musculoskeletal function across the sprawling–parasagittal transition and can ultimately be used to test hypotheses of limb morphofunctional transformation in the synapsid fossil record.

## Data accessibility

Interactive versions of the figures can be accessed through a Shiny app in R; details of running the app are provided in the main supplementary information file. The MMA and ROM results from the Maya and OpenSim modelling are provided as a supplementary data file.

## Authors’ contributions

R.J.B. and S.E.P. conceived and designed the study. P.F-L. and S.R. collected the original muscle data. R.J.B. built and analyzed the musculoskeletal models. R.J.B. drafted the manuscript in consultation with S.E.P. All authors edited the manuscript and gave final approval for publication.

## Acknowledgements

We would like to thank Rachel Norris and Anthony Wilkes (University of Adelaide) for donating the echidna specimens; Emma Hanslowe, Jillian Josimovich, Bryan Falk, and Robert Reed (United States Geological Survey Daniel Beard Center) for donating the tegu specimens; and Tom French (Massachusetts Division of Fisheries and Wildlife) for donating the opossum specimens. We would also like to thank Ken Angielczyk (Field Museum) for continued advice and Peter Bishop (Harvard University) for providing constructive feedback on this work.

## Supplementary Information

Supplementary information includes table S1 with specimen staining and scanning parameters, supplementary figures S1-S3 with details of model construction and coordinate systems, and supplementary figures S4-5 with alternative plots of the results highlighting specific MMA comparisons.

We also include a description of how to run the accompanying R Shiny app that displays the results as interactive 3D plots.

